# Role of the membrane cholesterol-glycosphingolipid complex as a ‘transistor’ to regulate GSL receptor function and signaling of both lipids

**DOI:** 10.1101/137612

**Authors:** Anton Novak, Clifford Lingwood

## Abstract

Cholesterol and glycosphingolipids (GSL) are the major species that accumulate in plasma membrane lipid rafts. These complexes imbue the membrane with increased order, which in turn, plays a central role in the transmembrane signaling foci lipid rafts provide. In addition, both GSL and cholesterol binding can mediate (separate) signal pathways. We have shown that cholesterol and GSLs however, form a complex in which the GSL sugar is reoriented from a membrane perpendicular to parallel format, becoming largely unavailable for exogenous ligand binding. Similarly, the steroid hydroxyl is masked, restricting access of cholesterol ligands. This was observed in model and cell membranes and in human tumour frozen tissue sections. We now show the order of exogenous ligand binding plays a significant role to determine the extent of GSL or cholesterol receptor activity. Ligand binding to cholesterol enhances subsequent GSL recognition and vice versa, suggesting that ligand binding to “free” receptor (membrane perpendicular GSL carbohydrate, nonmasked cholesterol) can result in partial dissociation of the GSL/cholesterol complex to allow additional GSL ligand and cholesterol ligand binding. Since many GSLs can complex with membrane cholesterol, the binding of a single cholesterol ligand may unmask cholesterol-complexed GSL for increased binding of both a single or multiple GSL-specific ligands. We show that multiple cholesterol-masked GSLs can be coincident in tissues. This provides a mechanism for GSL-dependent signal amplification and diversification, representing a biological ‘transistor’, regulating amplitude and potentially, diversity of GSL signaling. The process represents a new mechanism of ‘cross-talk’ between GSL and cholesterol signaling. This is of clinical importance since we have found cholesterol/GSL masking applies to monoclonal anti GSL antibodies in development and in current use as antineoplastic therapeutics.

## Introduction

GSLs are involved in many signal transduction pathways[1], both directly[2] and as modulators of the transduction of other signals[3,4]. Their increased concentration, together with cholesterol[5], in membrane lipids rafts[6] provide the appropriate location for modulating [7] these foci of transmembrane signaling[8].

While aberrant GSL metabolism is the direct cause of the lysosomal GSL storage diseases[9], GSLs play a major role in other human diseases, such that inhibition of their synthesis results in the amelioration of clinical and animal disease model symptoms[10- 13].

Errors in cholesterol homeostasis are widespread causes of disease [14-20]. Membrane cholesterol can transduce signals[21] and, via lipid rafts, modulate the activity of many trafficking[22,23]and signaling processes[24-26].

The sugar-lipid conjugate nature of GSL structure results in complex modulationof membrane GSL receptor function [27-30]. One of the best defined mechanisms for this modulation is the masking of membrane GSL expression within the cholesterol/GSL complex [31-34]. This cholesterol –induced conformational change in the GSL carbohydrate, from ligand available, membrane perpendicular (uncomplexed GSL), to ligand unavailable membrane parallel (cholesterol complexed) GSL conformers is dependent on an H-bond network between the steroid hydroxyl and the anomeric oxygen and amino function of the GSL [32]. Cholesterol masking is maximized under conditions of minimal membrane acyl chain mismatch[33]. Increased acyl chain mismatch promotes lipid rafts [35] so cholesterol masking of GSLs may be also modulated at the raft-membrane interface [7].

Due to the increased levels of cholesterol in tumours [36], GSL masking is a common feature observed in human tumour biopsies[34] and may be a means to escape immune surveillance, providing a rationale for cholesterol depletion in tumour immunotherapy.

Significantly, several Mabs approved and in development for human cancer treatment, turn out to target tumour GSLs [37-39]. In our studies, we found that several GSL antigens were cholesterol masked in the same location in tumour frozen sections (e.g. Gb_3_ and SSEA1 in colon cancer[34]). This led to our consideration of the effect of cholesterol masking of one GSL on the cholesterol masking of another.

From our previous study on the effect of sequential ligand binding to Gb_3_ and to cholesterol in prostate cancer sections[34], we now propose a model in which cholesterol masking may provide a switch (transistor) for GSL antigen exposure and hence, a network of ligand activated, GSL dependent signal transduction.

## Materials and Methods

### Tissue staining

Frozen tumor tissues were obtained from the Department of Pediatric Laboratory Medicine at this hospital and the Ontario Institute for Cancer Research Tumor Bank (Toronto, Ont, Canada). Cryosections (6 **μ**m) were dried and fixed with 4% paraformaldehyde in phosphate-buffered saline (PBS) for 10min then treated ±10 mM MCD in PBS for 30 min at room temperature (RT). Under these conditions, MCD preferentially extracts cholesterol [40] and we routinely observed negative section staining with filipin.

For brightfield microscopy of sections stained with horseradish peroxidase (,HRP) endogenous peroxidase blocker (Universal Block, KPL, Inc., Gaithersburg, MD) was added for 20 min, then samples were blocked with 1% normal goat serum (NGS) in PBS (1% NGS) for 20 min. Sections were treated with primary antibodies or VT1-B subunit. All reagents were diluted in 1% NGS, and all toxin/antibody reactions were for 30 min. VT1-B subunit (1 **μ**g/mL) was purified in our laboratory [41]. VTB incubation was followed by rabbit polyclonal anti-VT1-B-6869 [42]diluted 1:1000. Unituxin (dinutuximab anti-GD2 mAb, humanized chimeric mAb 14.18) was generously provided by United Therapeutics Inc NC, USA and Mab F77 was a gift from Dr M Greene, University of Pennsylvania.Pa. USA. Primary antibodies were used at 2μg/mL Secondary antibodies conjugated to HRP were applied (1:500): HRP-goat anti-rabbit IgG, HRP-goat anti-mouse IgG, or HRP-goat anti-human IgG (Bio-Rad, 172-1050). HRP-goat anti-mouse IgG (Bio-RAD, Hercules, CA, 170-6516). Sections were incubated for 5 min with peroxidase substrate (ImmPACT diaminobenzidine, Vector Laboratories, Burlingame, CA). Nuclei were weakly counterstained with aqueous Mayer’s hematoxylin (Vector Laboratories, Burlingame, CA). Slides were dehydrated in ethanol, cleared with xylene and mounted in Permount (Fisher, Ottawa, ON Canada). Images were recorded with a Nikon microscope (Eclipse E4000) using the Nikon ACT-1 software, version 2.70 and a Sony camera using Infinity Lumenara software, version 6.5.4. Pixel quantification was performed on digital images using Image J software.

For fluorescence microscopy of filipin-stained sections, filipin III (Sigma, St Louis, MO, F4767) was dissolved in dimethyl sulfoxide/PBS (1:4) (50 **μ**g/mL) and added to sections for 30 min. After rinsing, slides were mounted in aqueous fluorescent-mounting medium (Dako Cytomation, Glostrup, Denmark). Images were recorded with a Leica microscope (DMI 6000 B) using the LA SAF software, version 2.2.1.

For double staining, sections were stained with filipin and VTB /antibodies. When filipin was used first, it was followed by VTB, primary antibody, then HRP conjugated secondary antibodies. When VTB, antibody was reacted first, filipin was added after the HRP-conjugated secondary antibody, followed by incubation with the peroxidase substrate. All incubations were at RT. Slides were mounted in the aqueous fluorescent-mounting medium and recorded with the Leica microscope.

s

## Results and Discussion

### Cholesterol/GSL masking provides a basis for amplification of GSL signaling

Our results on the effect of sequential binding of Verotoxin B subunit (to Gb_3_ GSL) and filipin (to cholesterol) in prostate cancer biopsy frozen sections[34], have been reconstructed in figure 1A, and provide the basis of our ‘transistor’ model (figure 1B) for the regulation of GSL receptor function in membrane GSL signaling.

**Figure1.**
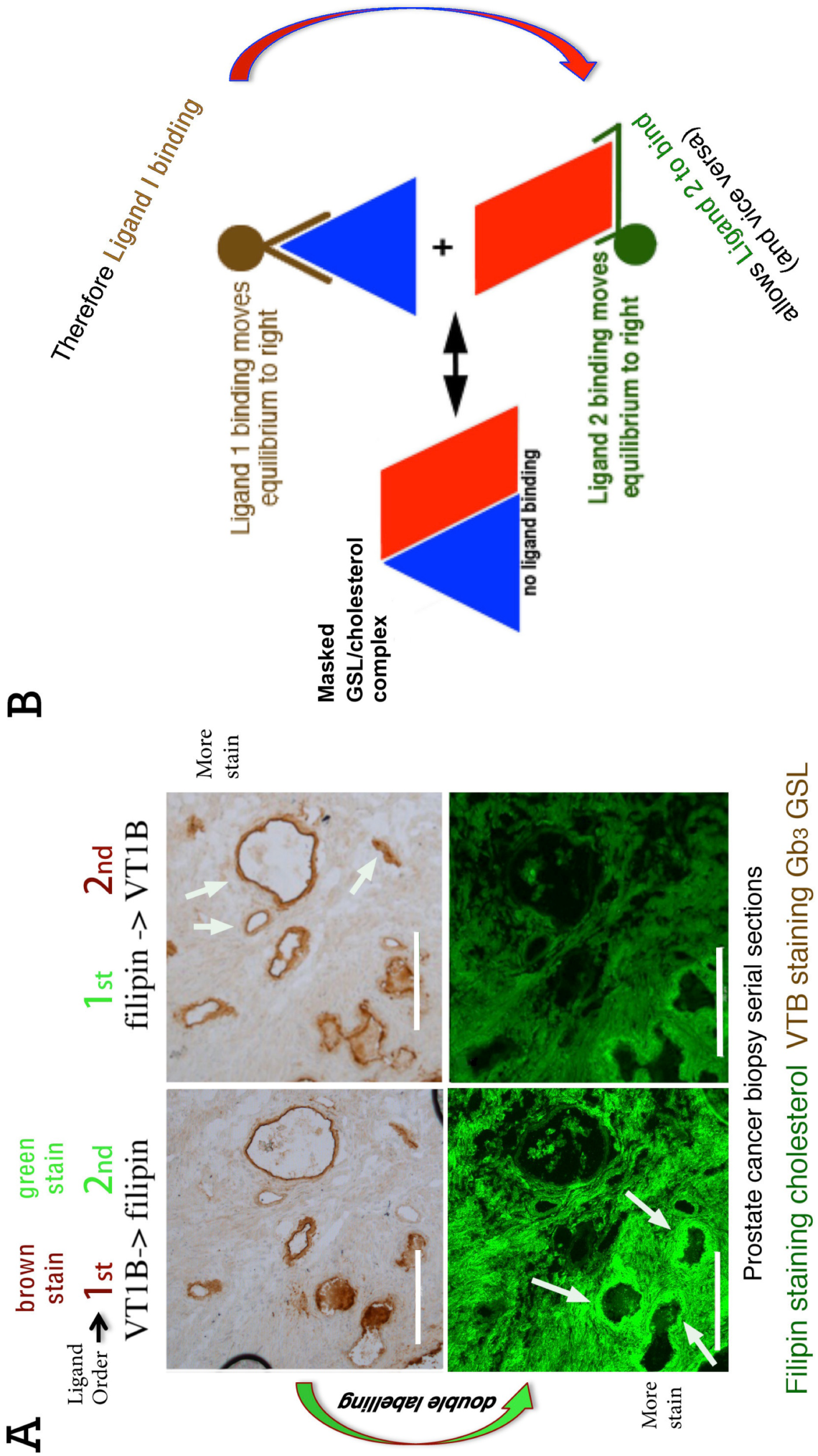
Double labeling of Gb3 and filipin to illustrate function of membrane GSL transistor. (**A**). Serial sections of a prostate cancer biopsy were co stained either with VTB/peroxidase (to detect Gb_3_, brown) or filipin-fluorescence (to detect cholesterol, green) or filipin first followed by VTB) [34]. Bar=200μm. Adding VTB first (top left) promoted subsequent filipin staining (bottom left). Filipin staining was most increased for vesicle showed the greatest initial VTB (Gb_3_) staining (lower left panel, arrows). In contrast, adding filipin first (top right) promoted VTB binding (not to all vesicles but see vesicles arrowed). (**B**) The explanation for this effect provides the basis of the GSL/cholesterol transistor. Binding of either ligand (VTB or filipin) to the small fraction of non-complexed lipid will result in a shift in the complexed vs. free equilibrium to the right, and thus dissociate a fraction of the complex, to promote the binding of the second ligand. This can be a switch or amplification for the second ligand binding.

An electrical transistor is a semiconductor which provides a switch mechanism for regulating current flow in an external circuit. A small current pulse through a transistor can induce the flow of a much larger pulse in the receiver (amplifier). The transistor can also allow a single electric input pulse to be split to many downstream receivers

Using serial prostate cancer frozen sections, a binding comparison was made when VTB or filipin was added first or second, and the detection of each ligand separately assessed and compared (fig 1A). The binding of filipin prior to VTB (right panels), resulted in increased detection of Gb_3_ by VTB compared to the same tubular structures in the tumour stained first with VTB (fig 1A top right, arrowed tubules). Not all VTB stained tubules show this amplification. Similarly, cholesterol detection with filipin was greater following initial VTB binding (left lower panel). Filipin staining was particularly elevated for tubules (fig 1 A lower panel left, arrows) not susceptible to filipin-increased VTB binding. This could be due to differential membrane cholesterol vs Gb_3_ concentrations.

Based on this data, we propose (figure1B) that ligand –receptor binding can partially unmask the cholesterol/GSL complex to amplify GSL or cholesterol mediated signaling pathways. In the GSL/cholesterol complex (represented by the red rhomboid, and blue triangle fig 1B), the availability of each lipid for ligand binding is restricted. This complex will, however, be in equilibrium with the free fraction of the membrane lipids. Ligand binding to either free lipid will drive the equilibrium a little to the right. This complex dissociation will provide additional unmasking of the other lipid to amplify binding of the second ligand. Thus, GSL binding increases the cholesterol available to bind an appropriate ligand (in this case, filipin). Such a ligand could contain a cholesterol binding motif[43,44], for example. Theoretically, due to the widespread nature of GSL masking by cholesterol[31,34], any GSL specific ligand[45] may unmask otherwise unavailable cholesterol. In the reverse case, a cholesterol binding ligand will similarly partially dissociate the GSL/cholesterol complex to partially unmask membrane GSL for increased ligand binding.

Our finding that the filipin cholesterol binding was increased more than VTB Gb_3_ binding is consistent with the proposal that more than one molecule of cholesterol is involved in the GSL/cholesterol complex[33] i.e. more cholesterol than Gb_3_ would be unmasked. This increased lipid receptor availability could take the form of allowing a lower concentration of the second ligand to bind the membrane lipid and would thus function as a ‘switch’, activated by the binding of the first ligand.

The signal amplification can be quantitated by ΔfreeGSL=k(nL2) or Δfreechol=k(nL1) where k is the coefficient of complex dissociation and L1 and L2 are the two ligands.

Since many GSLs are cholesterol masked[31], its is likely that any cholesterol binding species[44] could unmask a variety different GSLs if present, allowing diversification and/or amplification of a signaling route.

If two or more masked GSLs are masked within the same cholesterol domain, it is possible that, subsequent to GSL 1 ligand binding, a cholesterol binding ligand may then unmask both GSLs, resulting in signal diversification and the binding of one GSL ligand would result in the unmasking of other GSLs. This signal diversification would permit location dependent signal cross talk.

Cholesterol is also involved in many signal transduction pathways [21,46-48] and this model provides an interface between between GSL recognition and these pathways.

From analysis of figure 1A, we propose by analogy, that a biological ligand binding to a membrane GSL, which itself is either complexed, or not complexed to membrane cholesterol, can both amplify the signal transduced and split the signal into multiple downstream, GSL/cholesterol dependent signaling pathways. Thus, although by entirely different mechanisms, GSL/cholesterol signaling can therefore behave as a biological equivalent of a transistor.

### Mab anti tumour GSLs are subject to cholesterol masking

To emphasize the clinical relevance, particularly in cancer, we have investigated the effect of cholesterol masking on the tumour binding of two antiGSL Mabs developed for the treatment of human cancers.

**A) Mab F77.**This Mab was raised by immunization with a human prostate cancer cell line[37]. Prostate cancer is the cancer in which cholesterol plays a major role[49,50]. This antibody binds a H-blood group related carbohydrate expressed on GSLs and mucin[51]. Figure 2 shows the binding of Mab F77 to serial frozen tissue sections from a primary human prostate cancer biopsy, before and after beta-methyl cyclodextrin mediated cholesterol extraction. Mab F77 staining to the same tissue components was increased by 350% following MCD cholesterol extraction(primarily tubular structures [52]).

**Figure 2.**
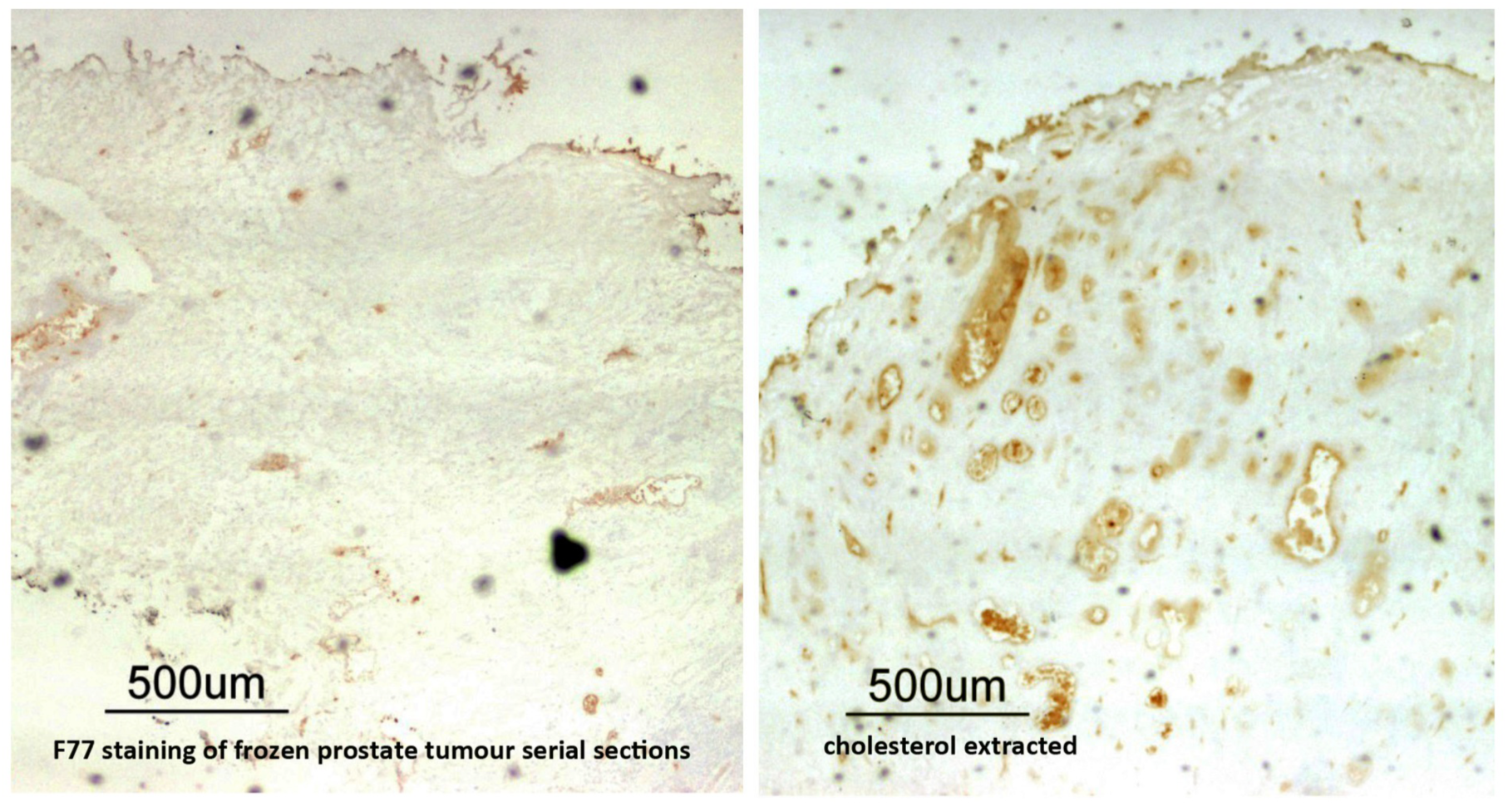
F77 Mab binding to prostate tumour is markedly increased by MCD cholesterol depletion. F77 binding to serial prostate cancer biopsy before and after cholesterol depletion. A few tubular structures are labeled before but many more after MCD extraction.

**B) Mab anti GD2 (Unituxin)** is the first FDA approved Mab for pediatric cancer (neuroblastoma [39,53]). Binding of this humanized mouse anti GD2 ganglioside antibody was increased to156% following cholesterol depletion, (figure 3). This significant increase in tumour binding offers the opportunity to increase the clinical efficacy of this antineoplastic approach. In this regard, it is of interest to note that a cyclodextrin based method to deplete membrane cholesterol in pediatric patients has been developed for Niemann-Pick disease [54], which is without apparent long term side effects.

**Figure 3.**
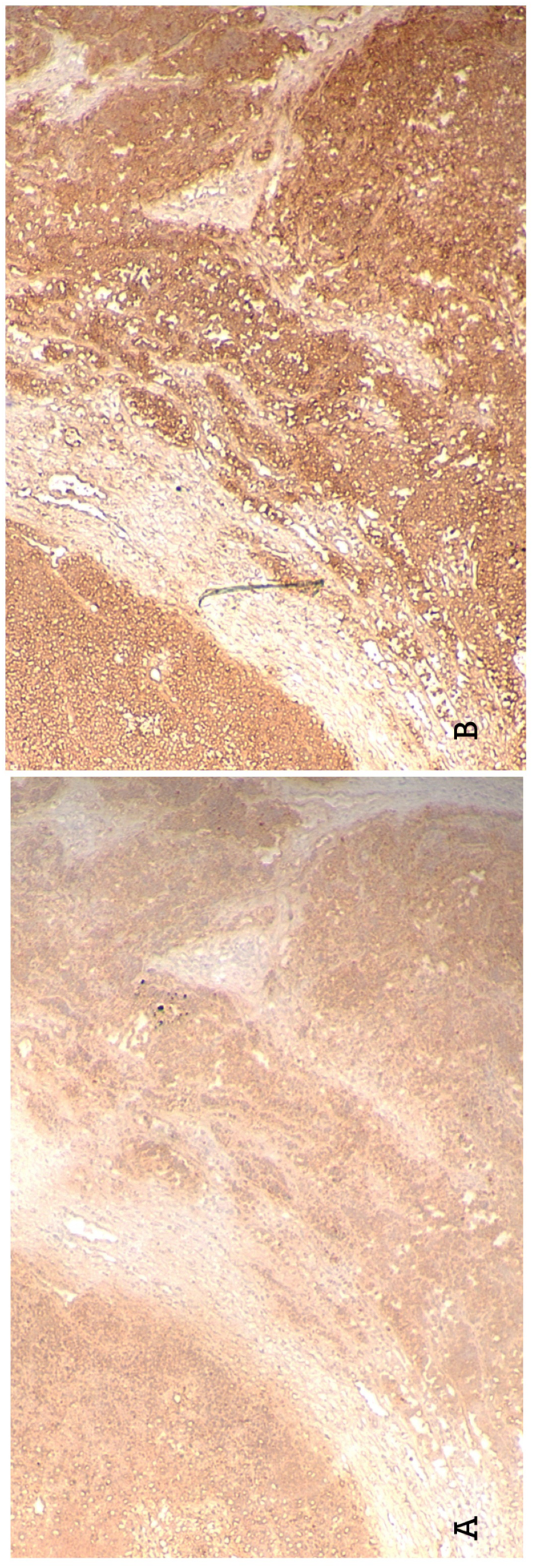
AntiGD2 staining of pediatric neuroblastoma. Anti GD2 (Unituxin) showed extensive staining of neuroblastoma sections (left panels) but cholesterol extraction resulted in a significant increase in tumour reactivity (right panels). Bar=250μm.

### Coincident unmasking of different GSL antigens

Our transistor model implies that ligand binding to one masked GSL could result in unmasking of a second masked GSL if it were in the same location. We therefore investigated whether two GSL could be masked in the same location. Figure 4 shows the localization of unmasked GD2 and Gb_3_ in colon cancer serial sections. MCD untreated sections show weak GD2 staining within the tumour. Cholesterol depletion markedly unmasks GD2 which is largely uniformly distributed within the tumour, but is not found in the columnar epithelial cells (fig 4B*). Gb_3_ is not detectable in untreated sections, except in these epithelial cells (fig 4C*). Significant Gb_3_ is unmasked (fig 4D) which is more widely distributed and overlaps unmasked GD2 (fig 4B,D arrows). Non-coincidental GSL unmasking demonstrates the selectivity of cholesterol-based GSL masking (i.e. it is not a non-selective general enhancement, but rather depends on the presence of specific GSLs). Co-incidental unmasking of different GSLs indicates locations in which our transistor model of signal diversification could operate.

**Figure 4.**
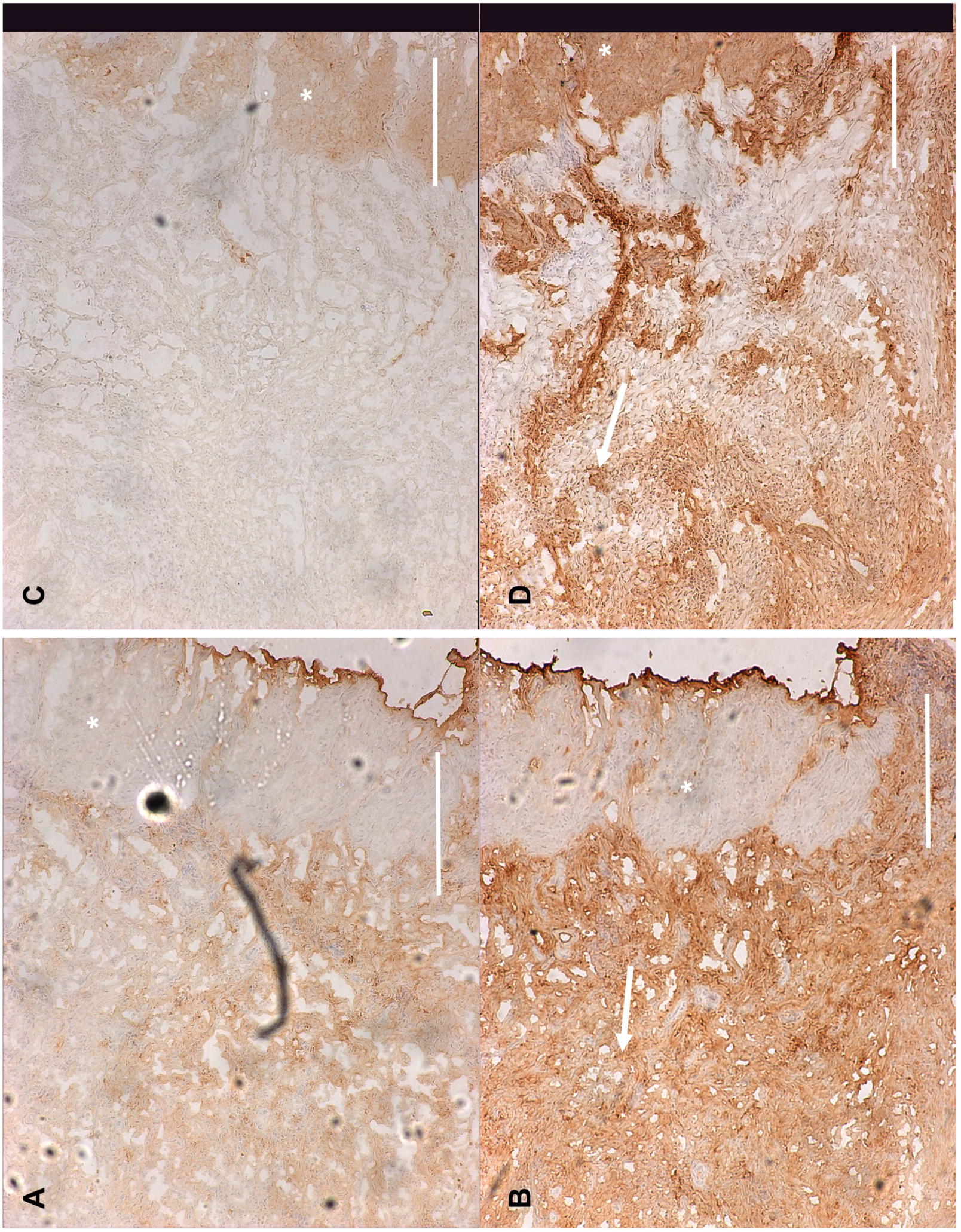
Colabelling of unmasked GD2 ganglioside and Gb_3_ in colon carcinoma biopsy sections. Serial human colon carcinoma frozen sections were stained for GD2 (A) or Gb3 (B) using Unituxin or VTB, prior to (A,B) or after (C,D), MCD cholesterol extraction. A marked increase is staining or these glycolipids within the bulk of the tumour was observed (arrows).* marks columnar epithelial cells. Inverted images are shown. Bar=50μm.

In breast tumour (figure 5), these antigens are, interestingly, partially colocalized within the tubular ducts of the tumour. GD2 is expressed within the tumour stroma and also within the outer layers of ducts (figure 5A). Increased GD2 expression is observed in both locations after cholesterol depletion (figure 5B). In contrast, Gb_3_ is expressed in a few blood vessels in the untreated section, but following cholesterol extraction, blood vessel staining is strongly increased, together with staining of the middle and inner layers of the duct structures. Thus, these unmasked GSL antigens overlap within the middle layer of the ducts but are distinct in the outer (GD2) and inner (Gb_3_) tumour duct layers.

**Figure 5.**
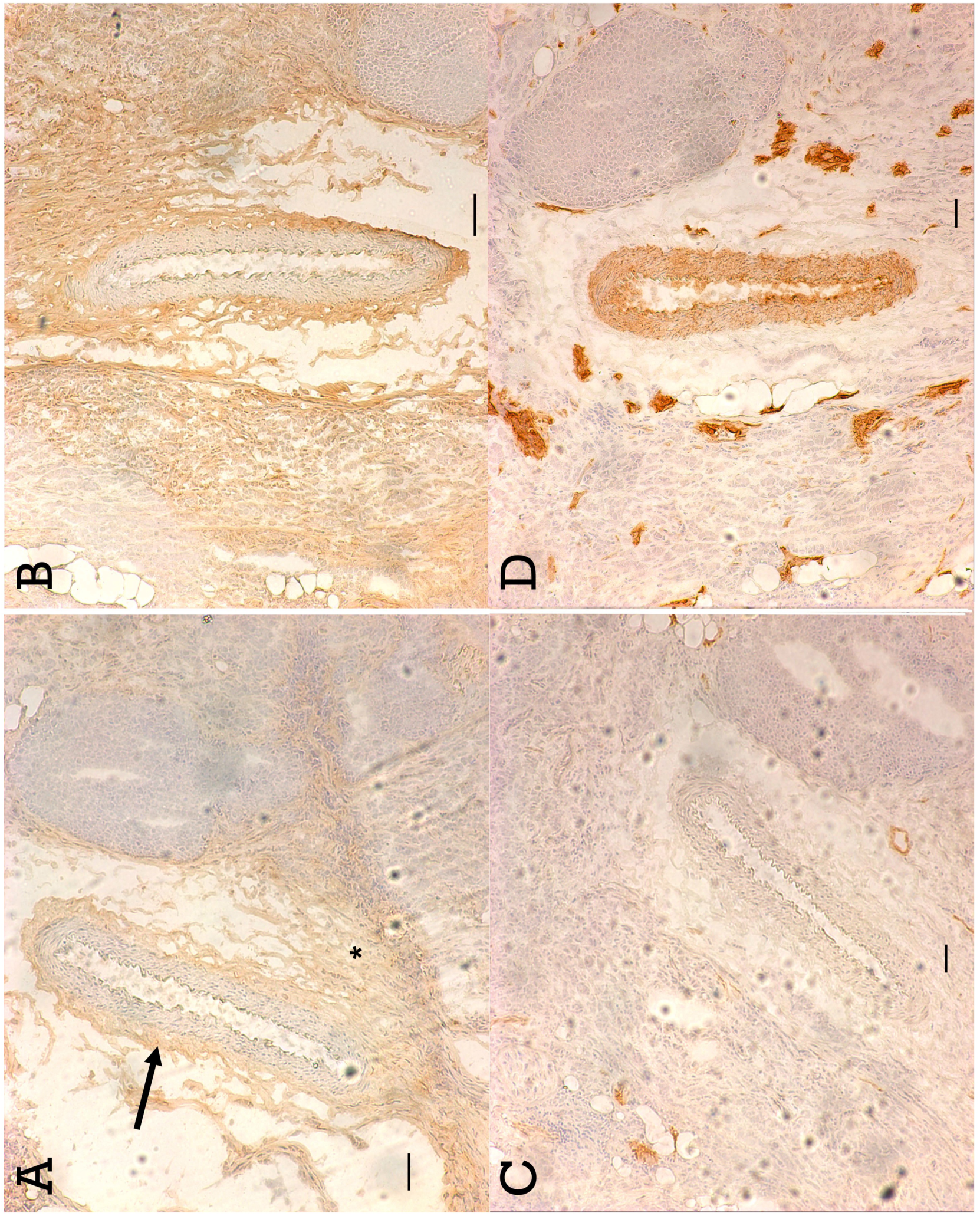
Colabelling of unmasked GD2 ganglioside and Gb_3_ in breast carcinoma biopsy sections. Serial human breast carcinoma frozen sections were stained for GD2 (A,B) or Gb3 (C,D) using Unituxin or VTB, prior to (A,C) or after (B,D), MCD cholesterol extraction. Little initial VTB blood vessel staining was markedly increased after cholesterol depletion and the central and luminal cells of tumour ducts were labeled. More of the tumour stromal tissue was initially antiGD2 labelled (A*) as were outer layer cells of tumour ducts(A arrow). Both were significantly increased by MCD cholesterol extraction. Bar=100μm.

Thus, a transistor based signal might be selectively propagated to the middle layer interface cells of the duct structures from either ligation of the outer GD2 masked layer or the inner Gb_3_ masked layer.

## Significance

Our studies demonstrate a new mechanism for the physicochemical regulation of cellular GSL and cholesterol receptor function, whereby molecules within a membrane lipid complex can ‘talk’ to each other. The transistor-like mechanism for signaling we have demonstrated in (what was) living tissue, can likely be mimicked in model membranes, since cholesterol masking of GSLs is readily apparent in such vesicles [31], opening the possibility that such a biological transistor might be useful in bridging biological and semiconductor based computation [55]. Biological computer prototypes have been made based on synthetic DNA[56] or RNA[57] but the present model would be the first ‘transistor’ operating in normal unmodified cells.

## Acknowledgement

This work was supported by a Hospital for Sick Children intramural grant from the Garron Foundation and by United Therapeutics.

